# Generation of apical-out nasal organoids to facilitate viral infection and drug screening

**DOI:** 10.1101/2025.10.07.680885

**Authors:** Georgios Stroulios, Mathieu Hubert, Wing Chang, Allen Eaves, Sharon Louis, Philipp Kramer, Caroline Tapparel, Salvatore Simmini

**Affiliations:** STEMCELL Technologies UK LTD, Cambridge, UK; Department of Medicine and Molecular Microbiology, University of Geneva Medical School, Geneva, Switzerland; STEMCELL Technologies Inc., Vancouver, BC, Canada; Terry Fox Laboratory, BC Cancer, Vancouver, BC, Canada

## Abstract

Advanced culture systems such as organoids can serve as powerful platforms to study epithelial physiology, as they recapitulate the organisation and many key functions of the tissue of origin. The nasal epithelium is the first respiratory epithelium that is exposed to inhaled airborne pathogens. As a result, it is crucial to model host-pathogen interactions occurring in this tissue. To facilitate the efficient modelling of these interactions, we have developed a method to generate *de novo* apical-out nasal organoids from nasal epithelial cell aggregates. Optimisation of this method revealed a stark tissue-specific effect of the culture temperature, as apical-out nasal organoids were generated in much higher efficiency at 32.5 ^°^C, compared to more widely used temperatures of 37°C. These organoids are composed of ciliated, basal and goblet cells and are produced in a completely standardised and scalable manner, devoid of any extracellular matrix hydrogel. Moreover, they displayed high homogeneity in size and cellular composition, as well as susceptibility to viral infections and capability to model antiviral drug responses. Here, we describe a method for the efficient and reproducible generation of apical-out nasal organoids with high potential to be utilised in host-pathogen interaction studies and personalised medicine from easy-to-access nasal swabs.

## Introduction

The nasal cavities form a complex organ that exerts crucial functions, which include olfactory sensing, as well as filtering and conditioning inhaled air. The nasal mucosa lines most of the cavities’ luminal surface, with an epithelium composed of columnar and cuboidal cells attached to a basement membrane. The anatomical location of the nasal mucosa at the uppermost part of the respiratory tract renders it an attractive site for putative pathogens to enter the host^1^. The exposure and susceptibility of the nasal epithelium to airborne pathogens present a substantial threat that can result in significant clinical burden.

A prime example of these pathogens is Respiratory Syncytial Virus (RSV), an airborne virus of great public health concern, since nearly every child is infected by RSV already prior to age 2 and reinfections occur regularly throughout life^2^. After acquisition, the virus mostly remains in the upper respiratory tract by infecting ciliated cells of the nasal epithelium but can progress to the lungs and cause severe lower respiratory tract infections (LRTIs) in immunocompromised patients, elderly individuals, young children and neonates^3^. To date, three vaccines (Arexvy, Abrysvo, mRESVIA) and two monoclonal antibodies (palivizumab, nirsevimab) exist to prevent RSV infection in elder patients and children, respectively, and more than 20 vaccines are in clinical stage of development^4^. In contrast, therapeutic options remain limited, with ribavirin as the only antiviral recommended to treat RSV-associated LRTIs, especially in immunocompromised patients^5–7^.

The nasal microenvironment plays a pivotal role in the entry and replication of RSV^8^, but also several other airborne viruses such as human and mammal-adapted avian influenza A viruses^9^ or the Severe Acute Respiratory Syndrome - coronavirus 2 (SARS-CoV-2), the virus responsible for the recent coronavirus disease 2019 (COVID-19) pandemic^10^. Since many more pathogens target the nasal mucosa, researchers have been eager to study the ontogeny and pathogenesis of infectious diseases in relevant model systems. To this end, a number of different *in vivo* models have been utilised to study host-pathogen interactions alongside the molecular mechanisms they employ. Yet, the presence of several anatomic and cellular interspecies differences, as well as the absence of crucial reflexes such as sneezing and coughing, have severely impacted the translatability of results to human disease^11^.

Numerous *in vitro* models allow for the manipulation and study of host-pathogen interactions, a critical step in understanding the mechanisms of respiratory illnesses. While simplified models, mostly based on immortalised cell lines, allow the study of specific disease aspects at the molecular level, they cannot recapitulate the tissue organisation, systemic response and mechanical forces that *in vivo* models can offer. Efforts are currently aiming at increasing the complexity of *in vitro* culture systems, in terms of both composition and structure, in order to maximise their physiological relevance.

*In vitro* culture of primary human nasal epithelial cells (hNECs) holds the potential to fulfil the need of increased complexity, and as a result researchers have focused on developing models based on this cellular source. However, expansion and differentiation of hNECs has been historically challenging, usually relying on a variety of both commercially proprietary and non-proprietary media optimised to culture human bronchial epithelial cells (hBECs)^12–16^. Different protocols have been developed to allow the generation of Air-Liquid Interface (ALI) cultures in cell culture inserts^12,17,18^, which largely resemble the *in vivo* tissue regarding the cellular composition^19^. Yet, the ALI culture system has inherent limitations, which include the limited scalability potential, a characteristic with paramount importance in the context of host-pathogen interaction studies and high-throughput drug screening.

Nasal 3D culture systems, such as organoids and spheroids, possess features to overcome some of these shortcomings. Nasal organoids have been successfully used to study cystic fibrosis^20–22^, SARS-CoV-2 infection^23^ and ciliary dyskinesia^24^. Similarly to other epithelial organoid models, nasal organoid culture systems require the support of matrigel or similar extracellular matrix (ECM) hydrogels. This induces the stereotypical polarity conformation presented by epithelial organoid when embedded in an ECM, where the apical surface of the epithelium faces inwards and is therefore not accessible from the external environment.

Recently, the generation of nasal 3D culture systems that expose the apical side of the epithelium to the environment has been described^15^. These spheroids were derived from non-adherent epithelial sheets obtained from nasal brushings and have the potential to be utilised in host-pathogen interaction studies. However, they pose significant drawbacks including the rather limited scalability stemming from the absence of expansion capabilities and the use of undefined components, such as bovine pituitary extract or feeder extracts used in the medium.

Here we optimised a workflow and culture conditions to generate a novel 3D apical-out nasal organoid culture system and confirmed its susceptibility to RSV, as well as the capacity to detect antiviral drug effects. Similarly to the apical-out organoid model that we previously generated from bronchial epithelial cells^25^, these nasal apical-out organoids demonstrate potential to overcome the shortcomings of previously described nasal cultures. Moreover, their polarity orientation offers the capacity to efficiently allow their use in host-pathogen interactions studies and other assays that require access to the apical surface of the nasal epithelium. Finally, these models can be obtained from nasal stem cells expanded from a patient’s own nasal swabs, offering new opportunities for personalised medicine.

## Methods

### Expansion of primary human nasal epithelial cells

Primary human nasal epithelial cells (hNECs) used in the study are from non-smoking, healthy donors and purchased from EPITHELIX SARL (cat # EP51AB). Initial cryopreserved hNECs were seeded as passage one (p1) into T25 cell culture tissue flasks in complete PneumaCult™-Ex Plus Medium prepared as per the manufacturer’s instructions (STEMCELL Technologies, catalogue # 05040) and incubated at 37°C and 5% CO2 with a full medium change every second day. Once cells reached 60–70% confluency they were dissociated using the Animal Component-Free (ACF) Cell Dissociation Kit (STEMCELL Technologies, catalogue # 05426) following the manufacturer’s instructions.

To assess the effect of temperature on the expansion potential of hNECs, cryopreserved cells at p2 were thawed and expanded for one passage at 37°C. Upon reaching confluency, cells were collected and seeded in PneumaCult™-Ex Plus either at 32.5°C or 37°C. Cells were serially passaged upon reaching confluency at each culture passage, until the cultures ceased expanding. Population Doublings (PD) were calculated using the following formula:

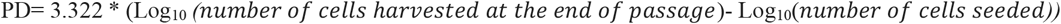

### Cultureware preparation

All plates used in organoid generation and in downstream assays were treated with Anti-Adherence Rinsing Solution (STEMCELL Technologies, catalogue # 07010) to reduce cell adhesion as previously described^25^. For all types of 24-well plates used, 500 μL of Anti-Adherence Rinsing Solution were added to each well. After centrifugation for 10 min at 1300 g, Anti-Adherence Rinsing Solution was removed and each well was washed once with 1 mL DMEM/F12 (STEMCELL Technologies, cat # 36254). The wells were used directly after removing the wash medium. Alternatively, 500 μL of fresh DMEM/F12 was added and they were stored for up to one week at 37°C.

### Generation of apical-out nasal organoids

Nasal aggregates were generated by first harvesting hNECs expanded in PneumaCult™-Ex Plus medium. Single cells were seeded in 24-well AggreWell™ 400 plates (STEMCELL Technologies, cat # 34411) in various concentrations. Complete PneumaCult™ Apical-Out Airway Organoid medium (AOAOM) was prepared as per the manufacturer’s instructions. Following seeding of hNECs, the AggreWell™ plate was centrifuged for 3 min at 100 g to sediment and aggregate the cells to the bottom of each microwell. Cultures grown at 37°C were generated with a density of 100 cells per microwell. Cultures that were incubated at 32.5°C had a density of 300 cells per microwell unless indicated otherwise. Similar to the donor optimisation required for apical-out airway organoids, the optimal aggregation time was determined by resuspending aggregates incubated in microwells in 24h intervals. The optimal timepoint was defined as the one where minimal fusion was observed at 24h post suspension. After the aggregates were generated and sufficiently matured, 1 mL fresh medium was added to each well. The aggregates were then resuspended using a P1000 pipette and each well was equally distributed to two wells of a 24-well plate previously treated with Anti-Adherence Rinsing Solution. Resulting aggregates were differentiated in suspension for a total of 15 to 28 days, with a 50% medium change performed every second day. Culture washing was performed by resuspending the culture using a P1000 pipette, allowing approximately one minute for the organoids to sediment to the bottom and subsequently removing 2/3 of the medium together with shed cells. Cultures were imaged using Leica DMi8. To quantify organoid generation efficiency, the total number of organoids generated by a single AggreWell™ well at day 28 was divided by the number of microwells, and this ratio was expressed as a percentage.

### Immunofluorescence staining

Terminally differentiated organoids were stained and imaged as described previously^25^. Briefly, apical-out organoids were fixed in Dents fixative (20% DMSO, 80% methanol) for 3h at room temperature with gentle shaking. Fixed organoids were permeabilised with 1% Triton X-100 (Sigma Aldrich, cat # 10789704001) in PBS, and blocked with 5% Normal Goat serum in PBS supplemented with 0.1% Tween-20 (Sigma Aldrich, cat # P9416) and 0.2% Triton X-100 (PBSTT). Primary antibodies were diluted in PBSTT and incubated with the cells overnight at room temperature in a tube with gentle agitation. Organoids were stained for ZO-1 (Invitrogen, cat #339100) and the epithelial markers acetylated α-TUBULIN (Sigma, cat # T7451), KERATIN 5 (Biolegend, cat # 905501) and MUC5AC (Abcam, cat # ab212636). Goat anti-Mouse IgG2b Cross-Adsorbed Secondary Antibody, Alexa Fluor 568 (Thermo Fisher Scientific, cat # A-21144), Goat anti-Mouse IgG1 Cross-Adsorbed Secondary Antibody, Alexa Fluor 488 (Thermo Fisher Scientific, cat # A-21121) and Donkey anti-Rabbit IgG (H + L), Alexa Fluor 647 (Thermo Fisher Scientific, cat # A-31573) were used respectively as secondary antibodies. Cells were washed with PBSTT and were incubated at room temperature with the respective secondary antibody in a tube with gentle agitation for 1h. Cells were washed again with PBSTT before staining with 4′, 6-diamidino-2-phenylindole (DAPI, Cayman Chemical, cat # 14285) and mounted using SunJin’s RapiClear 1.49 (Scientific Laboratory Supplies, cat # RC149001). Organoids were imaged using LEICA SP8 or DMi8. To assess the number of goblet cells, individual organoids stained with MUC5AC were imaged and counted manually.

### Dissociation of apical-out nasal organoids and ciliated cell counts

Apical-out nasal organoids were harvested either on day 15 or 28, transferred to a 15 ml tube and centrifuged at 150 g for 5 min and washed once with DMEM/F12. After an additional spin and removal of the supernatant, they were resuspended in TrypLE Express (Fisher Scientific, cat # 11558856) and incubated for 5-8 min at RT, before being dissociated to single cells by pipetting vigorously with a P1000 pipette. The single-cell suspension was diluted 1:1 with Trypan Blue (STEMCELL Technologies, cat # 07050), loaded to a hemocytometer and imaged in Leica DMi8, where the ciliated cells were counted manually.

### Organoid size, number and cilia presence homogeneity

Fully mature organoids were imaged using a Leica DMi8. Organoid size was measured by imaging mature organoid cultures and using Fiji^26^ to determine the Feret diameter. Organoid number and presence of beating cilia in organoids was manually determined using brightfield microscopy.

### Quantitative PCR

Total RNA was isolated from Ap-O NO using the Qiagen RNeasy Mini Kit (cat # 74106) following the manufacturer’s instructions. 500 ng of RNA was DNase treated (Invitrogen, cat # AM2222) as per the manufacturer’s protocol and then reverse-transcribed to cDNA using SuperScript III (Invitrogen, cat # 18080-044). TaqMan gene-specific assay primers and probes were obtained from Integrated DNA Technologies (Table 1) and reaction mixture from Applied Biosystems (cat # 4352042). Samples were amplified as follows: denaturation at 95 °C for 3 min followed by 40 cycles at 95 °C for 5 s and 60 °C for 30 s. The mRNA expression levels of cellular genes were normalised with that of *TBP*.

**Table 1:** Accession number of assays used to characterise apical-out nasal organoids.

| Gene | IDT assay |
| --- | --- |
| *KRT5* | Hs.PT.58.14446018 |
| *ITGA6* | Hs.PT.58.453862 |
| *FOXJ1* | Hs.PT.58.40371261 |
| *TUBB4B* | Hs.PT.58.15334509.g |
| *SCGB1A1* | Hs.PT.58.1190800 |
| *MUC5AC* | Hs.PT.58.24664713.g |
| *TSLP* | Hs.PT.58.22464960 |
| *FOXI1* | Hs.PT.58.40514934 |
| *TBP* | Hs.PT.39a.22214825(1) |

### RSV infection

Terminally differentiated organoids generated at 32.5°C from hNECs collected from three independent donors were harvested on day 28, washed once with complete PneumaCult™ AOAOM without heparin and infected with respiratory syncytial virus carrying a mCherry reporter gene (RSV-mCherry^27^) at MOI of 0.001 to 1 in the presence or absence of increasing doses of ribavirin (Merck, cat # R9644). At 24h post-infection, mCherry signal was measured using a BioTek Cytation 5 plate imager (Agilent) and quantified using the GEN5 software. The inhibitory concentrations (IC50) of ribavirin were determined for each donor using GraphPad Prism 10 software.

## Results

### Generation of Apical-Out Nasal Organoids Using the Airway Organoid Protocol

Given the biological similarities between hBECs and hNECs, we tested whether PneumaCult™ Apical-Out Airway Organoid Medium (AOAOM), a formulation previously optimised for the generation of apical-out airway organoids (Ap-O AO)^25^, had the potential to support the generation of apical-out nasal organoids (Ap-O NO) (Supplementary Figure 1A). Single hNECs were aggregated in AggreWell™ 400 micropatterned plates to promote the generation of homogenous-sized Ap-O NO. Large numbers of similar-sized hNECs aggregates (Supplementary Figure 1B) characterised by a dense, spherical morphology with minimal protrusions (Supplementary Figure 1C) were efficiently established across multiple donors by day 15. However, at this stage a few aggregates survived and successfully differentiated into organoids with a dense central core and ciliated cells on the outer surface of the epithelium. Most organoids exhibited fragile and irregular morphologies with numerous shed single cells, indicating extensive cell death due to suboptimal culture conditions (Supplementary Figures 1D-E). The observed irregular morphology strongly suggested suboptimal culture conditions, as dense, spherical structures with minimal protrusions has been previously associated with successful and well-differentiated Ap-O AO^25^. To benchmark the efficiency of Ap-O NO generation, we compared our results with historical data of Ap-O AO^25^ (Supplementary Figure 1F). This analysis confirmed that Ap-O NO were generated with an efficiency of approximately 10%, much lower than the 60% efficiency observed in Ap-O AO derived from cells at the same passage. Fluorescent immunostaining analysis confirmed that these organoids were composed of acetylated α-TUBULIN-expressing ciliated cells, KERATIN5 (KRT5)-positive basal and MUCIN 5AC (MUC5AC)-positive goblet cells (Supplementary Figure 1G).

These results indicate that, similarly to the airway, functional nasal epithelial organoids can be generated with an apical-out configuration. Yet, the poor morphology and the rapid deterioration of the cultures, in addition to the high cell death, resulted in an overall low organoid yield that strongly indicated the requirement for optimisation of the whole workflow.

### Physiological Temperature Enhances the Generation of Human Apical-Out Nasal Organoids

*In vivo*, the human nasal mucosal epithelium is exposed to a wide range of different internal and external environmental temperatures. In nasopharyngeal regions, temperatures were measured between 29°C and 37°C (average 32.6°C) with an ambient temperature of 23°C^28^. Nasal epithelial temperatures were found to correlate with ambient temperatures, with cold air reducing epithelial temperature^28^. Since cultures of different hNEC model systems are routinely generated and maintained at 37°C, we assessed whether different temperatures could influence the biological properties of nasal cells *in vitro*. To do so, we compared both the expansion and differentiation properties of cells cultured at 37°C and 32.5°C, which was selected as the temperature measured most frequently at the nasopharyngeal region^28^. To assess temperature’s effect on differentiation, we first expanded all cells at 37°C. Subsequently, we initiated differentiation towards Ap-O NO at both 32.5°C and 37°C, using a protocol similar to the one established for airway organoids (Figure 1A). Aggregates were successfully generated at 32.5°C and when transferred into suspension culture exhibited a similar morphology to those observed at 37°C, characterised by dense, mostly circular structures with a defined border that occasionally displayed irregular protrusions on the outer surface (Figure 1B). This morphology was largely preserved in suspension cultures until day 15, although a few rare protruding bubble-like structures could be observed (Figure 1C). Importantly, at 32.5°C cultures could be maintained for at least 28 days (Figure 1D) with minimal morphological changes and cell shedding throughout the duration of the differentiation, in contrast to the high cell death and declining morphology observed in cultures at 37°C (Supplementary Figures 1D-E). The differences in robustness were more apparent when we measured the generation efficiency of organoids derived from the two different temperatures (Figure 1E). Cultures maintained at 32.5°C demonstrated a higher organoid generation efficiency at day 15 when compared to those generated at 37°C (average organoid number per well of 669 at 32.5°C and 361 at 37°C). The improved capacity of organoids at 32.5°C to preserve their morphology for longer periods was also translated into higher organoid survival rates. Cultures at 32.5°C maintained the same number of organoids until day 28, by which time organoids generated at 37°C had completely dissociated (average organoid number per well of 627 at 32.5°C and 0 at 37°C). At that timepoint, organoids generated at 32.5°C displayed a halo of beating cilia on the outer surface, a feature similar to that observed on day 15 in cultures generated at 37°C.

**Figure 1:**
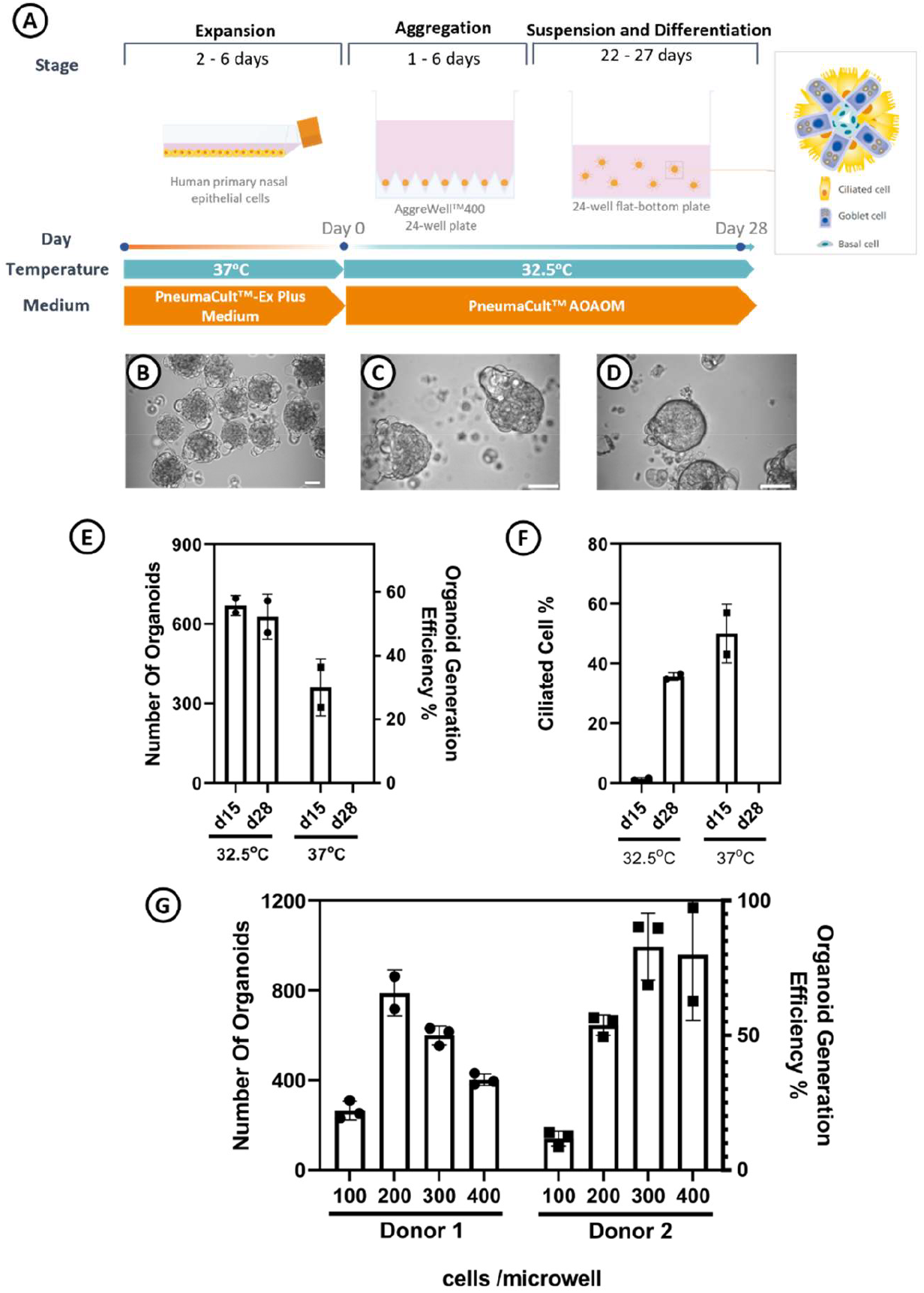
Lower culture temperature and higher seeding density improve Ap-O NO yield. A. Schematic of the modified protocol indicating the different temperatures used in each stage. B-D. Representative brightfield image of cultures generated at 32.5°C on day 1 (B), day 15 (C), and day 28 (D). E-F. Bar graphs comparing organoid number (E) and percentage of ciliated cells (F) in cultures generated at 32.5°C and 37°C at days 15 and 28 respectively. Points represent technical replicates. (n=1 donor) G. Graph depicting the changes in organoid number and generation efficiency with modification of the seeding density in two donors at 32.5°C at day 28. Points represent technical replicates. (Scale bars = 50 μm)

To determine the differentiation potential of hNECs in these two temperatures, we compared the percentage of cells that displayed beating cilia on day 15 and day 28. On day 15, cultures at 37°C displayed robust ciliogenesis based on counts of ciliated cells (average 49%), whereas those at 32.5°C had few ciliated cells (average 1.4%). However, organoids maintained until day 28 at 32.5°C demonstrated much higher numbers of ciliated cells (average 35.1%) that more closely resemble physiological levels^29^ (Figure 1F).

Finally, we optimised the workflow by testing different cell aggregation densities in AggreWell™ 400. We have previously described that 100 hBECs per microwell were sufficient to efficiently generate Ap-O AO and that higher seeding densities caused excessive cell shedding during suspension, resulting in organoids of similar size at the end of the workflow^25^. To determine if phenotype and generation efficiency of Ap-O NO were also affected by the initial seeding density, we seeded 100, 200, 300 and 400 cells per microwell of an AggreWell™ 400 plate at 32.5°C. Interestingly, all tested seeding densities were able to generate Ap-O NO, but higher seeding densities demonstrated an increased organoid generation efficiency (Figure 1G). Organoid forming efficiency appeared to plateau between 200 and 300 cells seeded per microwell, with increasingly high seeding densities having a detrimental effect on the final organoid counts. As a result, we defined 300 cells per microwell as the optimal seeding density to be used for the efficient generation of Ap-O NO.

### Validation of the Optimised Workflow for Robust Generation of Ap-O NO

To evaluate our refined workflow, we derived Ap-O NO from 3 distinct donors and assessed several key aspects, including organoid generation efficiency, phenotype and cellular composition. Considering the differences observed in organoids differentiated at 32.5°C, 37°C, and across different timepoints, characterisation and comparison analysis were performed at the timepoint where each temperature condition demonstrated its optimal phenotypic results. Therefore, we compared Ap-O NO generated at 37°C on day 15 (non-optimised workflow) to organoids generated at 32.5°C on day 28 (optimised workflow). The optimised workflow resulted in a significantly higher number of organoids and greater generation efficiency for all 3 hNEC donors, compared to the non-optimised workflow. For the optimised method, organoid counts averaged 949 (Donor 1), 788 (Donor 2) and 1030 (Donor 3), with corresponding organoid generation efficiencies of 79%, 65.6%, and 85.8%. In contrast, the non-optimised method yielded average organoid counts of 498 (Donor 1), 248 (Donor 2) and 835 (Donor 3) and efficiencies of 41.5%, 20.6% and 69.5%, with Donors 1 and 2 displaying highly significant differences between the two conditions (Figure 2A). Similar morphological improvements were observed in cultures generated from either of the two new donors cultured at 32.5°C, when compared to 37°C. This indicated that the efficiency observed at lower temperature is an inherent characteristic of the modified approach, rather than stemming from a distinctive potential of the specific donor.

**Figure 2:**
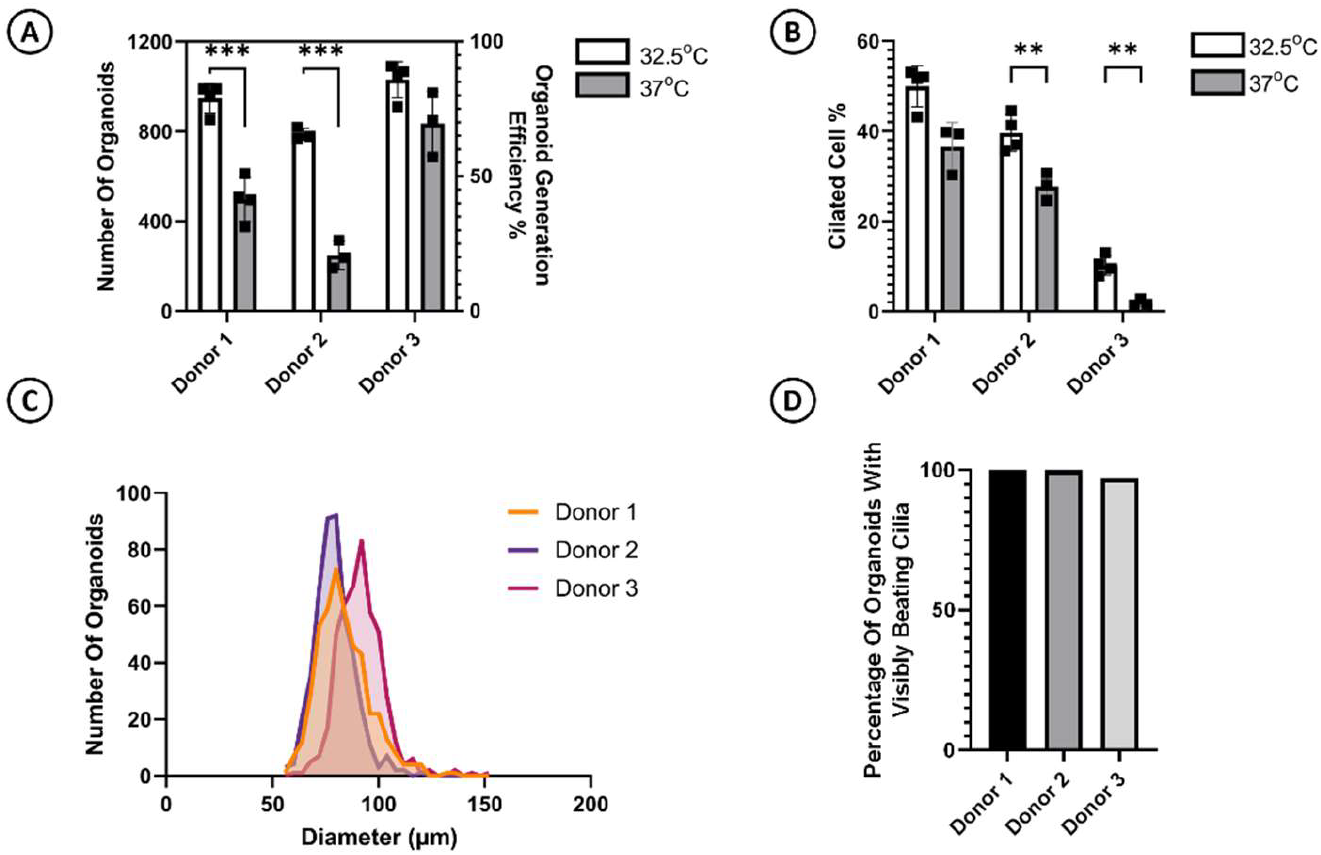
Modified workflow supports high yield, uniformity, and ciliary function in Ap-O NO. A-B. Comparison of the optimised and non-optimised protocols across three donors, showing quantification of organoid number (A) and ciliated cell percentage (B). Measurements were taken at day 28 for the optimised protocol and day 15 for the non-optimised. Points indicate technical replicates. C. Frequency distribution of Feret diameters for p4 organoids from three donors, based on measurements of at least 450 organoids per donor. D. Percentage of organoids with active ciliary beating observed on the outer surface, assessed from 200 organoids per donor.

All Ap-O NO across conditions and donors were characterised by functional cilia that could be observed beating on the outer side, confirming the apical-out polarity. Quantification of the ciliated cell percentage indicated that the differentiation potential of Ap-O NO generated at 32.5°C was increased in all donors, with Donors 2 (39.62%) and 3 (10.2%) showing a significant increase compared to what measured in cultures from the same donors at 37°C (Donor 2 27.76%, Donor 3 1.92%). Interestingly, Donor 3 overall displayed a noticeably reduced population of cells with beating cilia compared to the other donors at both temperatures, highlighting the capacity of the model to preserve inter-donor heterogeneity *in vitro* (Figure 2B).

The use of the AggreWell™ platform to generate aggregates of homogeneous size has previously resulted in the establishment of Ap-O AO with a high degree of size homogeneity^25^. To determine whether this characteristic was also preserved in Ap-O NO, we generated cultures using p3 hNECs and measured the Feret diameter of the resulting organoids (Figure 2C). This analysis demonstrated that Ap-O NO derived from all the tested donors displayed a remarkable size homogeneity, with an average diameter (+ standard deviation) of 83.49±12.2 μm, 79.16±9.08 μm and 91.37±10.72 μm for Donors 1, 2 and 3 respectively. The coefficient of variation was calculated as 14.63% for Donor 1, 11.48% for Donor 2 and 11.74% for Donor 3, further highlighting the uniformity of the culture. This homogeneity was further confirmed by the quantification of ciliogenesis, as the vast majority of organoids (97%) derived from all 3 donors displayed visibly beating cilia at the outer surface (Figure 2D).

Since PneumaCult™-Ex Plus could support the proliferation and serial passaging of hNECs, we tested whether these cells were also capable of retaining their differentiation potential at late passages. We harvested hNECs at the end of p4, the last passage they displayed proliferation, and used them to generate organoids using our e optimised workflow. Even at the latest passage, the organoid generation efficiency remained above 50% across all donors, with each culture well capable of producing more than 600 organoids per AggreWell™ well of a 24 well plate (Supplementary Figure 2A). When compared to the results obtained using cells of lower passages, the differentiation potential of high-passage hNECs seemed to be reduced (Supplementary Figure 2B) but still remained within physiologically relevant levels.

Finally, we characterised the cellular composition of Ap-O NO generated with our optimised protocol by harvesting them at day 28 and assessing the RNA levels of common markers of differentiated cell types (Figure 3A). When compared to the RNA profile of hNECs cultured in PneumaCult™-Ex Plus expansion medium, basal cell markers ITGA6 and KRT5 were significantly downregulated in Ap-O NO. In contrast, ciliated cell markers FOXJ1 and TUBB4B were significantly upregulated as a result of ciliated cell differentiation. Upregulation of goblet cell marker MUC5AC was also detected in two out of three donors. Presence of other secretory cells, such as club cells, could not be confirmed based on their marker (SCGB1A1) expression. Upregulation of markers characteristic of other rare cell types such as tuft (TSLP) and ionocytes (FOXI1) were not detected. Interestingly, the neuroendocrine cell marker ASCL1 was found to be upregulated, albeit the relatively high Ct values (>36) do not allow for a definitive conclusion of the presence of this cell type. Immunocytochemical stains of fixed Ap-O NO detected a robust apical border on the outer surface, characterised by expression of tight junction protein ZO-1 (Figure 3B). Moreover, further analysis confirmed the presence of differentiated cell types. The ciliated cell marker acetylated α-TUBULIN was detected in all donors, with the localisation of cilia on the outer surface further confirming the apical-out polarity of the organoids (Figure 3C). MUC5AC-positive goblet cells could also be readily detected (Figure 3D), as were KRT5-positive basal cells (Figures 3C, 3D). To better understand the frequency and distribution of goblet cells, we assessed the presence of MUC5AC-positive goblet cells in at least 170 individual Ap-O NO per donor (Figure 3E). Most (77.39%, 72.15% and 95.41% for Donors 1, 2 and 3 respectively) of the organoids contained at least 2 goblet cells, with a few (16.95%, 21.59% and 2.29% for donors 1, 2 and 3 respectively) containing a single goblet cell. Ap-O NO devoid of goblet cells were rarely observed (5.65%, 6.25% and 2.29% for donors 1, 2 and 3).

**Figure 3:**
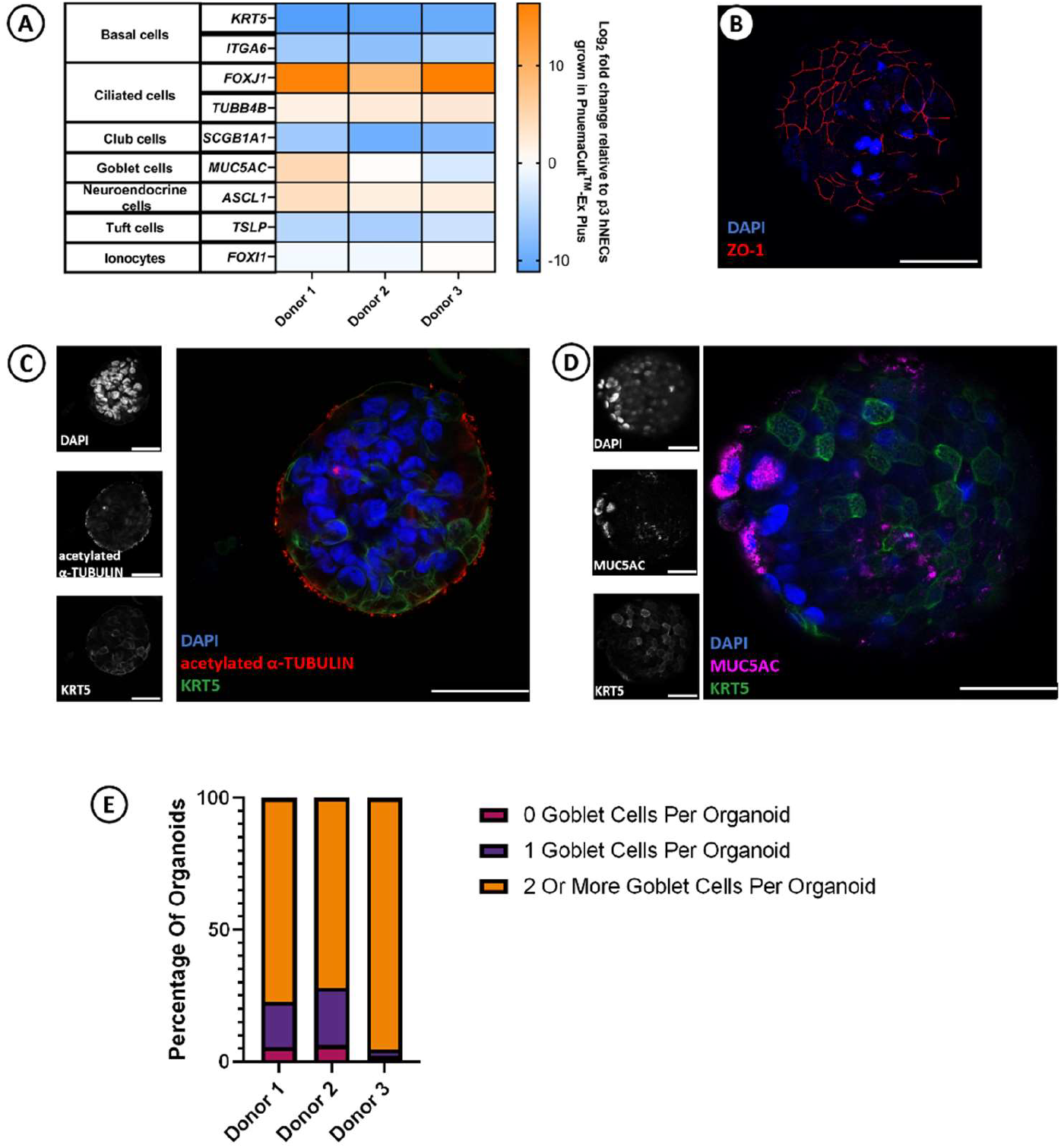
Successful differentiation and expression of key lung cell markers with the improved Ap-O NO workflow. A. Fold changes in the RNA levels of key lung differentiation markers in p4 hNEC-derived organoids from three donors. Expression was normalized to p4 hNECs cultured in PneumaCult™-Ex Plus. B-D. Immunofluorescence staining of organoids for the tight junction marker ZO-1 (red, B), the ciliary marker acetylated α-tubulin (red, C), the goblet cell marker MUC5AC (magenta, D), and the basal cell marker KRT5 (green, C, D). Nuclei are counterstained with DAPI (blue). E. Stacked bar graph highlighting the percentage of organoids that contain 0, 1 and at least 2 goblet cells per organoid for each donor.

Collectively, these data validate our optimised workflow for supporting an efficient and robust differentiation of Ap-O NO from hNECs in an ECM-free manner across multiple donors. Such Ap-O NO contain multiple differentiated cell types representative of the nasal epithelium *in vivo* and display a high degree of size homogeneity, making them well suited for multiple downstream high throughput assays.

### Lower Culture Temperature Impairs hNEC Proliferation

We next investigated the effect of temperature on hNEC proliferation by culturing hNECs derived from 2 donors in PneumaCult™-Ex Plus expansion medium at either 37°C or 32.5°C. We then compared cellular morphology and population doublings across multiple passages. The morphology of hNECs cultures at 37°C was characterised by the presence of small, tightly packed cells in the first 3 passages (Supplementary Figure 3A). A morphological decline was observed by p4, characterised by the formation of cell clumps with a “bubbly” appearance. Signs of stress were evident, with cells displaying an abnormally enlarged cytoplasm and vacuoles (Supplementary Figure 3B), leading to growth arrest at p5. Similar to cultures grown at 37°C, cells cultured at 32.5°C exhibited a tightly packed, good morphology during the first passage that declined at day 9 when cultures ended (Supplemental Figure 3C). Comparison analysis of the respective population doublings (PD) confirmed that hNECs cultured at 32.5°C were proliferating less (average PD until growth arrest 3.74) and slower (required 9 days to reach confluency), than cells cultured at 37°C (average PD until growth arrest 6.76 and required between 4 to 7 days to reach confluency) (Supplemental Figure 3D). These results clearly indicated that temperatures lower than 37°C could not support an efficient expansion of hNECs *in vitro*.

### Impact of Culture Temperature on Ap-O AO Generation

To assess whether modifications related to the temperature of the culture were tissue-specific, we tested if 32.5°C had any effect on the generation and morphology of Ap-O AO. After expansion of hBECs (n = 2 at p5) at 37°C, single cells were seeded in AggreWell™ 400 plates, at a density of 100 cells per microwell in PneumaCult™ AOAOM, either at 37°C or 32.5°C. At the end of the workflow on day 15, aggregates differentiated at 32.5°C in both donors had largely fused and formed one big aggregate of irregular shape (Supplementary Figure 4A). These structures displayed protruding cells, suggesting the presence of extensive apoptosis (Supplementary Figure 4B). Moreover, beating ciliated cells were not detected on the epithelial surface at this timepoint, implying that the lower temperature could have severely impacted the differentiation capacity of the hBECs. In contrast, cultures incubated at 37°C resulted in well-defined Ap-O AO and limited fusion, indicating the donor-specific optimisation time to reduce fusion events was successful (Supplementary Figure 4C). In addition, these organoids displayed the expected morphology, with a dense core and mostly spherical shape, with a clear darker border and minimal protrusions (Supplementary Figure 4D). Ciliated cells could be readily identified beating, indicating the presence of robust differentiation.

Collectively, these observations suggest that the reduction in temperature, which had a paramount effect in the generation efficiency and morphology of apical-out nasal organoids, had adverse effects in apical-out airway organoid generation. As a result, the beneficial effects of apical-out organoid generation at 32.5°C proved to be nasal-specific.

### Functional Assessment Of Ap-O NO For Viral Infection And Antiviral Testing

Similar to the airway, the nasal epithelium functions as a barrier to the outer environment and is targeted by a multitude of airborne pathogens. To test if Ap-O NO would be susceptible to viral infections and verify their potential to model these interactions, we infected terminally differentiated organoids with RSV. Exposing the organoids to increasing multiplicities of infection (MOI) of a recombinant RSV carrying an mCherry reporter (RSC-mCherry)^27^, allowed us to quantify the mCherry signal as a surrogate of infection. At 24 hours post infection (hpi), we observed a correlation between the viral inoculum and the RSV-mCherry area, which confirmed the susceptibility of Ap-O NO to RSV (Figure 4A). To test the capacity of the organoids to model antiviral drug effects, we next performed a dose-response assay with ribavirin, administered 2 hours before infection (Figure 4B). Ap-O NO treated with 75 μM ribavirin showed markedly reduced mCherry signal compared to control, indicating a high antiviral efficacy in pre-treatment conditions (Figure 4C). A dose-depended effect of Ribavirin was detected, with an IC50 of 15.54 μM (Figure 4D). Interestingly, the 3 different tested donors demonstrated varying IC50 values, ranging from 1 to 20µM (Figure 4E). Taken together, these results show that Ap-O NO are sensitive to RSV infection and can be used for drug discovery.

**Figure 4.**
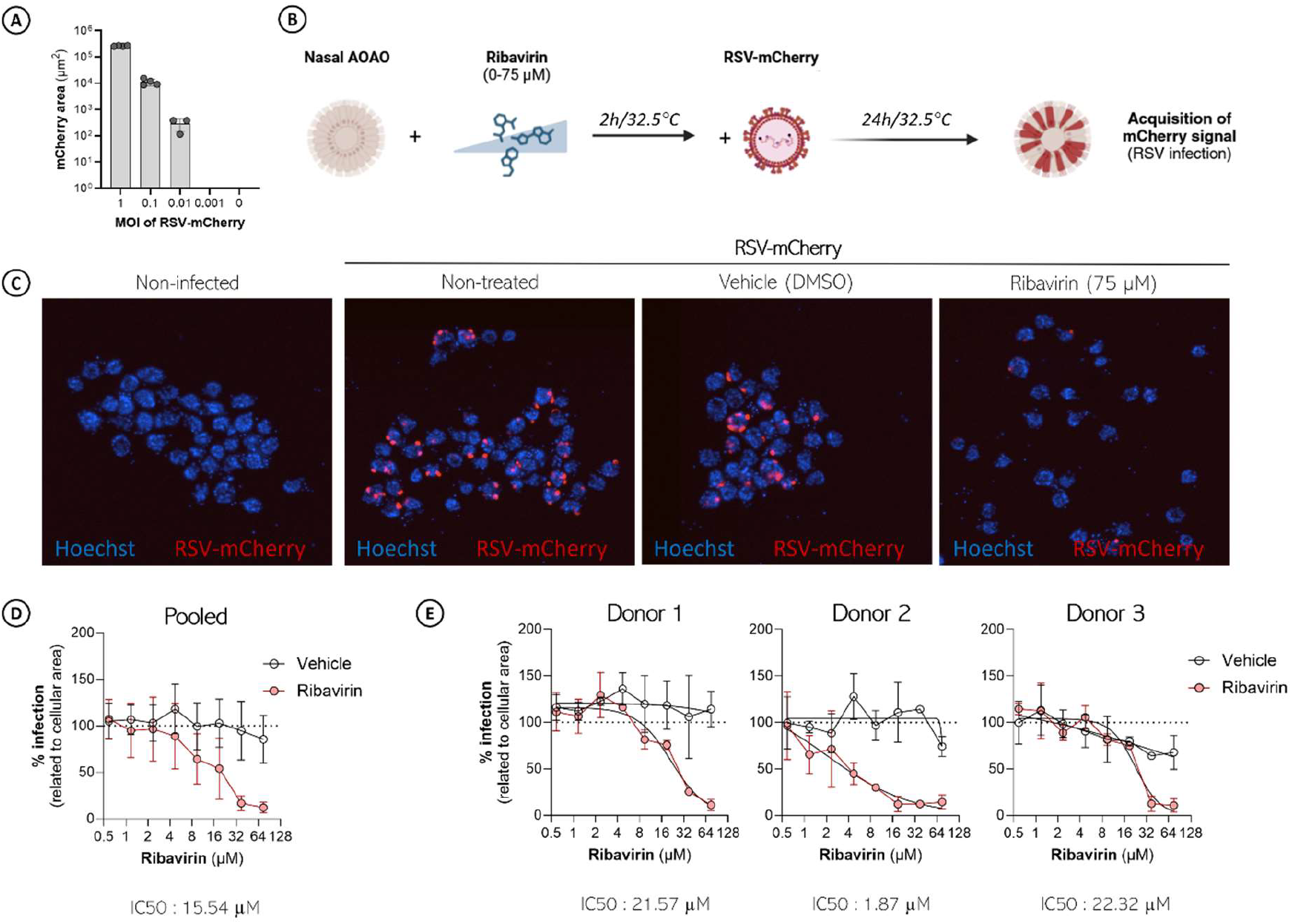
Ap-O NO are susceptible to RSV infection and are suitable for antiviral activity assessment. A. Representation of the dose-dependency of the mCherry area (expressed in µm^2^) in Ap-O NO in response to RSV-mCherry inoculum. Terminally differentiated Ap-O NO were exposed to RSV-mCherry at several multiplicities of infection (0.001 to 1) and the total mCherry area was measured as a surrogate of infection using a Cytation 5 plate imager. B. Terminally differentiated Ap-O NO were pre-treated with increasing concentrations of ribavirin (0 to 75 µg/mL) or vehicle (DMSO) alone. After incubation for 2h at 32.5°C, treated Ap-NO were exposed to RSV-mCherry (MOI 0.1). At 24h post-infection, mCherry signal was monitored by fluorescent microscopy as a surrogate of infection. C. Representative examples of micrographs taken from non-infected and RSV-infected Ap-O NO (MOI 0.1) in the absence or the presence of 75µM of ribavirin or vehicle alone. Nuclei are stained in blue (Hoechst) and RSV-infected cells are shown in red (mCherry). D. Dose-response analysis of inhibition of RSV-mCherry (MOI 0.1) by ribavirin (red) or vehicle (white) in Ap-O NO collected from three independent donors. Data are presented as mean±standard deviation (SD) of the three individual donors. The IC50 value is indicated below the graph. E. Individual dose-response analysis of each donor represented in D. Data are presented as mean ± standard deviation (SD) of triplicates from one experiment. The IC50 values are indicated below each graph.

## Discussion

Here, we describe a new method for the generation of apical-out nasal organoids from primary human epithelial cells in the absence of any extracellular matrix. We detail the optimisation of several aspects of the workflow to efficiently generate and differentiate Ap-O NO from 2D-expanded hNECs. Differentiated Ap-O NO were composed of ciliated cells that presented beating cilia on the outer surface of the organoid, as well as basal cells and mucin-producing goblet cells. Moreover, Ap-O NO displayed high size homogeneity and demonstrated great potential to be used in host-pathogen interaction studies based on their susceptibility to RSV and capacity to model the response to antiviral drugs like ribavirin.

The low organoid generation efficiency observed in the non-optimised workflow could be a result of insufficient growth factor supplementation, as media developed for the bronchial epithelia were used to culture hNECs. Despite the phenotypic, structural and functional similarities shared by the two tissues, some differences might still exist at a molecular level. These could stem from the different germ layer of origin, as airway epithelial cells derive from the endoderm and nasal from the ectoderm^30,31^. Differences have also been described at the expression level and activity of several key transcription factors between nasal and airway epithelial samples^32^. Therefore, media optimised for other regions of the airway epithelium may inadequately represent the molecular niche of the nasal epithelium. For example, hNECs can be expanded in 2D using airway-specific media, however the colony forming efficiency drops dramatically at early passages^33^. This is something that is also observed in expanding hBECs, albeit loss of proliferation capacity is observed at much later passages. This indicates that the current media formulations for the expansion of airway cells can be further optimised to better recapitulate the nasal stem cell niche found *in vivo*. Similar differences can also be observed in differentiation assays. Like hBECs, nasal cells can be differentiated in air-liquid interphase using porous membranes and commercially available airway media^34^. However, the levels of ciliogenesis observed in these cultures supported by defined or undefined (containing bovine pituitary extract or calf serum) airway media formulations have been considered rather poor^35^. Further characterisation of the properties of the nasal epithelium both *in vitro* and *in vivo* will expand our knowledge regarding its physiology, resulting in the generation of novel *in vitro* models that more accurately recapitulate the *in vivo* tissue.

Interestingly, lowering the incubation temperature during differentiation improved organoid’s phenotype and generation efficiency of Ap-O NO. Extending the differentiation duration by two weeks promotes the development of well-differentiated Ap-O NO and optimises the utilisation of each donor’s cells, a significant advantage considering the relatively low expansion potential of hNECs *in vitro*. The presence of multiple differentiated cellular types, along with the observation of functionally beating cilia, indicate the functional resemblance of these organoids to the *in vivo* nasal epithelium.

It has been shown that temperature also influences pathogen replication in nasal cavity: lower temperatures enhance viral replication, while higher temperatures inhibit it^36,37^. Considering this viral temperature sensitivity has been described to be exploited by the host’s immune system^38^, it would be interesting to study its effects *in vitro*. Interestingly, similar effects are also detected in mammalian cells, as culture temperature higher than that observed *in vivo* was found to have similar confining effects in the proliferation of pluripotent cells^39^. Moreover, temperature was described to influence the differentiation potential of keratinocytes^40,41^, dermal cells that *in vivo* are exposed to similar temperature conditions as the nasal mucosa. Here we showed that lowering the culture temperature to 32.5°C during differentiation drastically improved both the efficiency and morphology of the resulting Ap-O NO, across different passages. These might serve as an indication that culture conditions such as temperature should be assessed in a tissue- and assay-specific basis to supplement or augment the effect of different factors used to mimic the molecular niche.

*In vitro* nasal models have been employed to study the effect and infection route of several pathogens, including Influenza A^9,42^, Rhinovirus 16^43^, RSV^34,44,45^ and SARS-CoV-2^10,45^. Interestingly, RSV infection was recently shown to exhibit higher replication and cytopathogenicity in infant-derived human nasal epithelial cultures compared to adult-derived counterparts^8^. Here, we demonstrated that Ap-O NO are suitable for antiviral drug discovery, and their scalability potential has promising applications in high throughput screenings. As a culture system, Ap-O NO present similar benefits to Ap-O AO, including a completely ECM-free and standardised workflow. However, the differences in the tissue accessibility offer additional advantages. Generation of both primary airway and nasal epithelial models requires a supply of donor cells, which have a finite lifespan. This results in a constant requirement for additional donors. Sourcing bronchial airway cells can be challenging, as they are routinely derived from patient materials such as lavages, resections and biopsies^46^. Yet, such invasive procedures are not commonly undertaken, especially in healthy individuals. Moreover, the properties of the cells like the morphology, differentiation potential and response to pathogens might be altered due to the individual’s disease background. Post-mortem tissues are often presented as an alternative, although the potential impact that death had on the cells’ molecular mechanism and the effect of processing at different timepoints have not been assessed. As a potential alternative, nasal cells can be obtained from both adults and children, with minimal invasiveness through nasal brushing. This ease of sampling gives researchers the opportunity to recruit large datasets with desired characteristics (for example age, sex, health status) to exclude potential bias arising from a small sampling group due to limited tissue availability. In addition, it allows the sequential sampling over multiple timepoints, enabling the study of phenomena in different age ranges, while maintaining the same genetic background. However, we think that researchers need to remain mindful of the differences between the nasal and airway epithelia to exclude potential generalisations.

The advantages of incorporating a diverse genetic background during drug screening was apparent in our testing, as different donors had alternate degrees of response to the same drug, ribavirin. Better understanding of the diverse drug actions that are influenced by the genetic background is something that is not easily feasible with immortalised cell lines due to their limited genetic diversity. Yet, organoid systems present not only an increased level of cellular and architectural complexity, but also the ability to reveal demographic differences in drug efficacy. Recently, Ap-O AO were utilised in RSV antibody neutralisation assays and described a cost-effective approach to not only detect neutralising antibodies directed to the RSV fusion (F) protein like immortalised lines exclusively do, but also those directed to glycoprotein (G)^47^. The characteristics of the Ap-O NO described here offer similar benefits and now present a valuable alternative to classical cell culture models. Furthermore, assessing the capacity of the system to capture the effect of different physiological tissue states (e.g., healthy, inflamed, chronic disease) has the potential to further increase its value. Leveraging the greater relevance of organoid systems and embracing the genetic differences between donors could lead to the development of treatments that are better tailored to an individual’s genetic background and a departure from generic treatments.

In summary, we have shown that apical-out nasal organoids can be successfully generated using an ECM- and chemically defined medium and workflow, initially developed for the airway and subsequently optimised for nasal tissues. These organoids closely resemble functional, cellular and architectural characteristics of the native nasal epithelium, exhibiting a high level of homogeneity and coordinated ciliary activity. Furthermore, their susceptibility to RSV infection and their dose-dependent response to ribavirin administration highlights their potential as platform for disease modelling and antiviral drug screening applications.

## Acknowledgements

The authors thank the Cambridge Advanced Imaging Centre for their support and assistance in this work. This research was funded by the Horizon 2020 grant OrganoVIR 812673 on the project ‘Organoids for Virus Research - An innovative training-ITN programme’. This research was also funded by STEMCELL Technologies Ltd., Cambridge UK and the Department of Medicine and Molecular Microbiology, University of Geneva Medical School, Geneva, Switzerland.

## Authors contributions

G.S., M.H., C.T. and S.S. designed the study. G.S. and M.H. performed the experiments. A.E., P.K., S.L., W.C., provided critical input and materials. S.S. and K.P. provided guidance. G.S., M.H., C.T. and S.S. wrote the manuscript with contributions from all the authors.

